# The function of the prereplicative complex and SCF complex components during *Drosophila wing* development

**DOI:** 10.1101/219220

**Authors:** Hidetsugu Kohzaki

## Abstract

Chromosomal DNA replication machinery functions in the growing cells and organs in multicellular organisms. We previously demonstrated that its knockdown in several tissues of *Drosophila* led to a rough eye phenotype, the loss of bristles in the eye and female sterile. In this paper, we investigated in detail the wing phenotype using RNAi flies, and observed that the knockdown not only of Mcm10 but also of some other prereplicative complex components demonstrated wing phenotypes, using Gal4-driver flies. Surprisingly, some SCF complex components, which control cell cycle progression via protein degradation, also showed the wing phenotype. These results showed that the DNA replication machinery contributes to wing development independent of growth, probably through defects in DNA replication and protein degradation at specific places and times.

**Summary statement:** We recently outlined the prereplicative complex components, including Mcm10 and SCF complex functions, during *Drosophila* wing development. In this paper, we detail these findings.

## Introduction

DNA replication machinery is essential for cellular growth. The origin recognition complex (ORC) is the platform for the DNA replication initiation complex. In *Drosophila*, Cdt1-Mcm2-7 is thought to be the licensing factor that directs the S phase for each cell cycle. Considerable evidence indicates that Mcm2-7 is also a replicative helicase (Masai et al., 2010; Hua and Orr-Weaver, 2017). The Cdc45, Mcm2-7, and GINS (CMG) complex is the eukaryotic DNA replication fork complex in the *Drosophila* embryo (Moyer et al., 2006). DNA polymerases, including Pol α-primase, Pol δ, and Pol ε, are the enzymes responsible for the elongation phase of DNA replication. The yeast Cdc6 is a loader of Mcm2-7 onto the DNA replication origin (Pacek et al., 2006; Borlado and Méndez, 2008). Though Cdt1 lacks enzymatic activity, and bears little resemblance to any other protein of known molecular function, it is essential for origin licensing in all eukaryotes (Pozo and Cook, 2017). The recent finding that Cdt1 has a second essential role in the cell cycle during mitosis underscores the importance of fully understanding its function (Varma, 2012). Recently, cryo-EM structures showed that each element of the Orc-Cdc6-Cdt1-Mcm2-7 (OCCM) intermediate plays a distinctive role in orchestrating the assembly of the pre-replication complex (pre-RC) by ORC-Cdc6 and Cdt1 (Zhai et al., 2017).

The activity of CDKs and other kinases at the G1–S and G2–M transitions must be tightly regulated to prevent inappropriate cell cycle progression (Reinhardt, and Yaffe, 2013). The SCF complex controls cell cycle progression by the degradation of proteins, including CycE, the CDK inhibitor p27^Kip1^, and so on (Lipkowitz and Weissman, 2011), after these kinases phosphorylate these substrates. This complex consists of Skp1–cullin 1 (Cul1)–F-box protein. The cell cycle is regulated by the SCF complex and anaphase-promoting complex/cyclosome (APC/C) multisubunit RING finger E3s. These complexes are targeted to specific substrates via interchangeable substrate recognition subunits, including F-box proteins for SCF and cell division cycle 20 (Cdc20) and Cdh1 for APC/C. Cul1 is an essential subunit of the SCF (Skp1, Cul1/Cdc53, F-box proteins) ubiquitin E3 ligase complex that targets many phosphorylated substrates, such as p27^Kip1^, IκB, β-catenin and Orc1, for κ ubiquitin-dependent degradation (Kipreos et al., 1996; Araki, et al., 2005). Although other cullin complexes are less well characterized, they all seem to function as ubiquitin E3 ligases by binding the RING finger proteins Roc1 or Roc2(RBX/HRT), which recruit the ubiquitin E2 conjugating enzymes for polyubiquitination. Furthermore, the silencing of Roc1a, one of three *Drosophila* Roc1 orthologues (Roc1a, Roc1b, and Roc2), was sufficient to suppress the disappearance of Cdt1 (the licensing factor complex subunit) after irradiation (Higa, 2003). Cdt1 is specifically polyubiquitinated by Cul4 complexes, and the interaction between Cdt1 and Cul4 is regulated in part by gamma-irradiation (Higa, 2003).

As noted above, the cell cycle progression needs growth and development (Kohzaki et al., 2018a).However, the retinoblastoma tumor suppressor protein (RB) has been shown to regulate cell cycle control by binding to E2F-DP1 (Nevins, 2001). The transcription factor, E2F-DP1 heterodimer, drives G1-S transition in the cell cycle. They express many proteins including Orc1which proceed cell cycles (Asano and wharton, 1999). It is now clear that the Rb/E2F pathway is critical in regulating the initiation of DNA replication (Asano, et al., 1996; Nevins, 2001).

In researching the availability of RNA interference (RNAi) in *Drosophila* (Hannon, 2003), we sought to compare the phenotypes of flies in which particular genes were knocked down in various tissues and stages (Kohzaki et al., 2018a). Previously, we knocked down DNA replication machinery using *Act5C-* and *tubulin-Gal4* drivers, which express target genes throughout the body. However, the knockdown of *Mcm2*, *Mcm4*, *Cdt1*, and *Cdc6* was lethal (Kohzaki et al., 2018a). Thus, in a further study, we used a tissue-specific RNAi knockdown system in combination with a Gal4-UAS system (Kohzaki and Murakami, 2018b). A transgene of interest, which is expressed with a Gal4-dependent promoter, is introduced into the embryo. By crossing with a fly expressing *Gal4* in a tissue-specific manner (*Gal4* driver), one can obtain flies that express the transgene in a tissue-specific manner. When a target gene’s antisense RNA is expressed using the tissue-specific *Gal4* driver system, the RNA transcribed from the target DNA forms double-stranded RNA, which can be destroyed by RNAi machinery, resulting in the depletion of the target gene product. We also observed whether Cul-4, dSkip-1/SkpA, Rbx1/Roc1, and Roc1b/Roc2 were lethal, and whether Cul-1 and Elongin C showed severe growth defects, when knocked down using *Act5C-* and *Tubulin-Gal4* drivers.

In *Drosophila* tissues, chromosomal abnormalities, including gene amplification and endoreplication, occur in a developmentally regulated manner (Claycomb et al., 2004). Recently, we showed that the DNA replication machinery required for *Drosophila* development and chromosome replication in the mitotic cell cycle is needed for gene amplification and endoreplication (Kohzaki et al., 2018a). Moreover, we reported that chromosomal DNA replication machinery plays an active role in tissue development in *Drosophila* eye development, and contributes to body development independent of growth (Kohzaki and Murakami, 2018b).

In this study, we first showed that the RNAi knockdown of chromosomal DNA replication machinery could induce abnormal wing formation. Next, we demonstrated that such knockdown not only affected DNA replication but also resulted in a rough wing phenotype, probably through defects in DNA replication. Finally, we revealed how the knockdown of DNA replication machinery, including Mcm10 and SCF complex components, led to novel phenotypes.

## Materials and methods

### Fly stocks

Fly stocks were maintained under standard conditions. The RNAi knockdown lines were obtained from the National Institute of Genetics (Mishima, Japan) and the Vienna Drosophila RNAi Center (Vienna, Austria). *Tubulin-p-Gal4*, *SD-Gal4*, *Vg-Gal4*, *en-Gal4*, *ptc-Gal4*, *dll-Gal4*, and *dpp-Gal4 (yw)*, were obtained from the Bloomington Drosophila Stock Center (Indiana, USA).

### Knockdown experiments

To investigate the function of DNA replication machinery in the rear of the wing imaginal disc, we knocked down the complex and SCF complex with *SD-Gal4* by means specific to tissue, time, and place (Kohzaki and Murakami, 2018b).

### Quantitative reverse transcriptase-PCR

Total RNA was isolated using Trizol Reagent (Invitrogen). Oligo dT primers and a Takara high fidelity RNA PCT kit (Takara, Kyoto, Japan) were used for generation of complementary DNA. Then, real-time PCR was performed using a SYBR Green I kit (Takara) and the Applied Biosystems 7500 real-time PCR system (Applied Biosystems, Foster City, CA, USA). RNA expression efficiencies decreased to 25% in every case (Kohzaki et al., 2018a).

### Stereo microscope and fluorescence microscope

Prepared specimens were examined using a fluorescence microscope (Olympus BX-50) equipped with a cooled CCD camera (Hamamatsu Photo ORCA-ER) and Aquacosmos image analysis software (Hamamatsu Photo ORCA-ER) (Kohzaki and Asano, 2016).

## Results

### DNA replication machineries, including Mcm10, are potentially involved in wing formation

We hypothesized that DNA replication machinery directly contributes to *wing* development. First, we performed the knockdown of DNA replication machinery using several Gal4 drivers for screening (Kohzaki et al., 2018a; Kohzaki et al., 2018b). Previously, we showed that the knockdown of Mcm10 by *tubulin-p-Gal4* and *SD-Gal4* led to wing formation defects (Kohzaki et al., 2018a). The knockdown of Mcm10 by wing-specific Gal4 and *SD-Gal4* showed wing defects, which were not observed with the use of *Vg* or e*n-Gal4* driver. We performed the knockdown of several DNA replication machineries, including Mcm3 and Pol *ε* 255KDa, using *SD-Gal4,* but we did *ε* not observe a resulting wing phenotype (Kohzaki et al., 2018a). These findings suggested that Mcm10 has a further function, in addition to its function in DNA replication.

In this study, we performed the knockdown of other DNA replication machineries, using *SD-Gal4.* In particular, we found that the knockdown of *Cdt1*, *Pol α-primase*, *RPA*, *Psf2* (a subunit of GINS), and *Rfc3* (an RFC complex) by *SD-Gal4* showed a wing phenotype. These results suggested that these factors, in particular, require the establishment of an elongation phase in DNA replication (Fig. 1). And the knockdown of Mcm10 by *SD-Gal4* showed wing defects, which were not observed with the use of *Vg*, *ptc*, *dll, dpp*, or e*n-Gal4* driver (Table 1). Though these results may have reflected defects in the unwinding stage, we do not, as yet, know the details. However, these findings suggest that DNA replication machineries, including Mcm10, have a function in wing formation, in addition to their function in DNA replication (Tables 1 and 2, and Fig. 2).

**Fig. 1.**
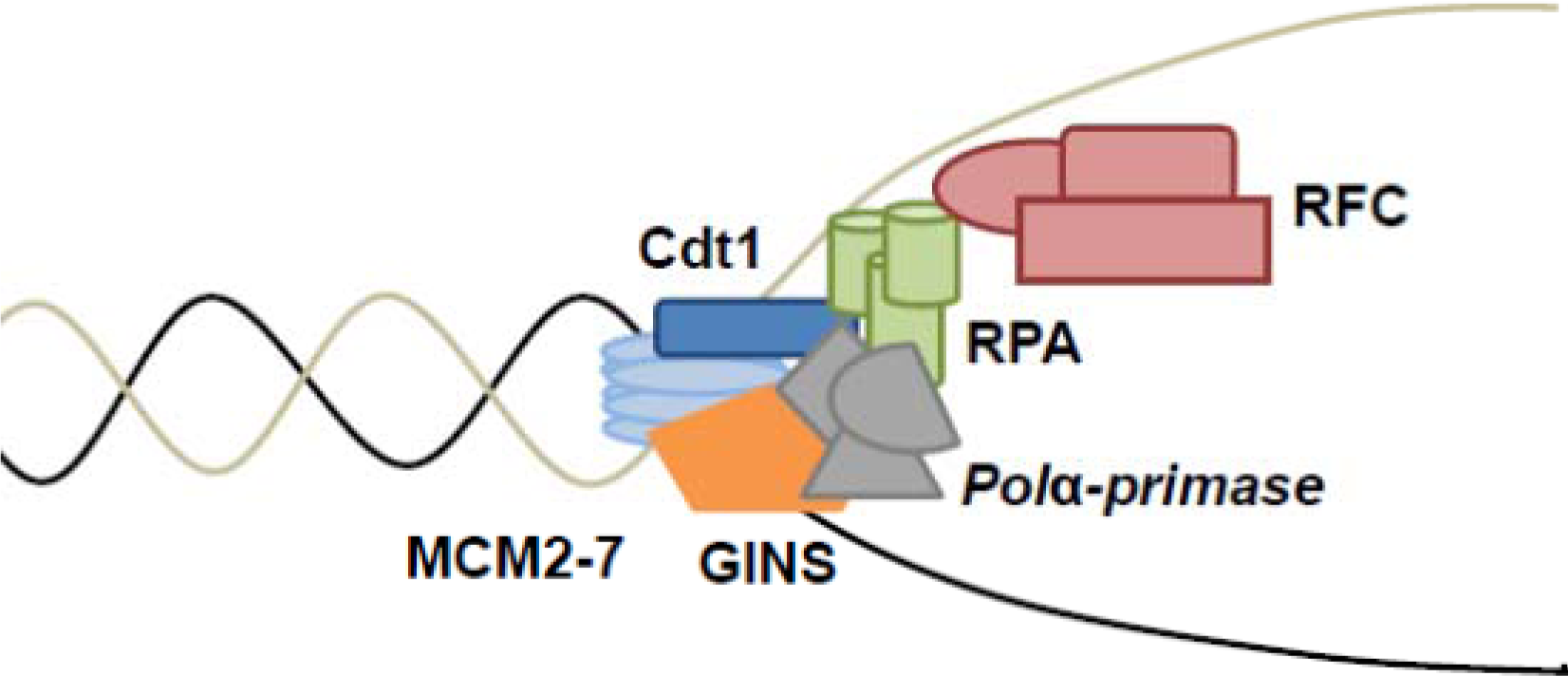
Creation of the elongation complex in DNA replication.

**Fig. 2.**
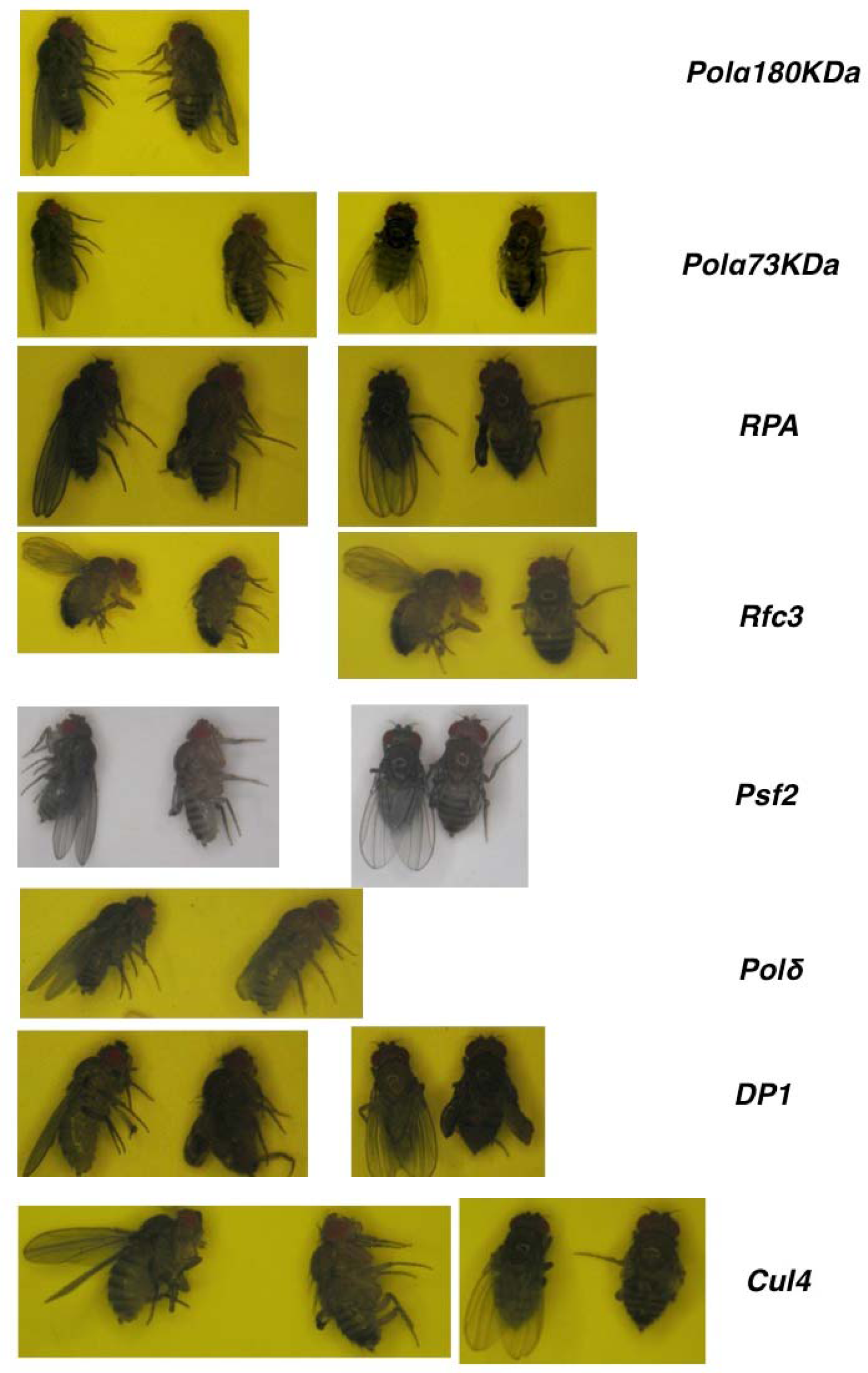
Knockdown of DNA replication machineries by *SD-Gal4* driver.

**Table 1.**
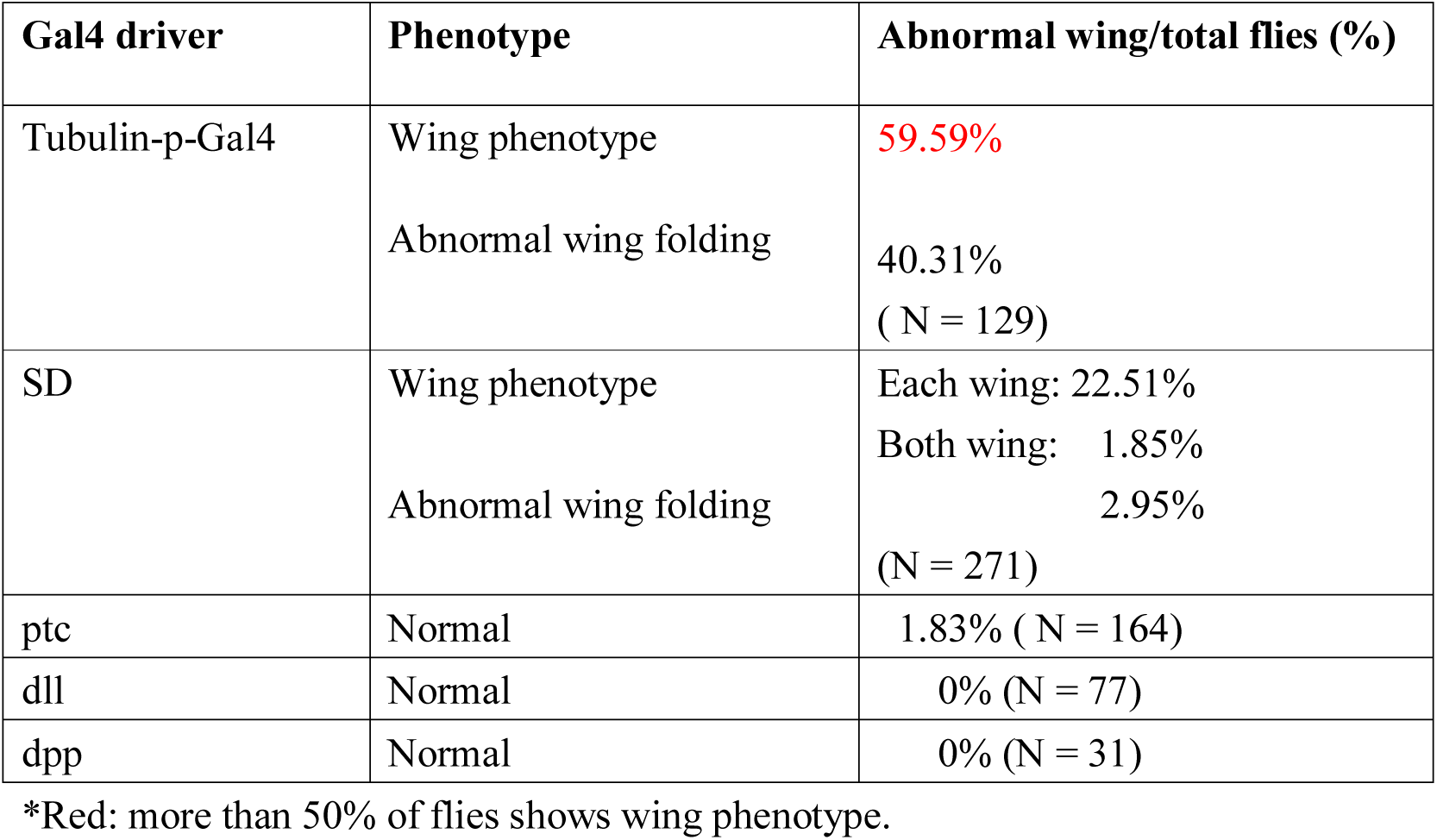
Summary of phenotypes induced by knockdown of Mcm10 with each Gal4 driver.

### The knockdown of the SCF complex by SD Gal4 drivers disturbed wing development

The degradation of proteins by the SCF complex and APC/C controls cell cycle progression at the G1–S and G2–M transitions (Kipreos et al., 1996; Lipkowitz and Weissman, 2011; Reinhardt and Yaffe, 2013). We observed that CycA, CycE, Cdc20, Cdh1, and Rca1 mutants were embryonic lethal. When knocking down Cul-1, Cul-4, dSkip-1/SkpA, Rbx1/Roc1, Roc1b/Roc2, and Elongin C using *Act5C* and *Tubulin-Gal4* drivers, we further observed whether Cul-4, dSkip-1/SkpA, Rbx1/Roc1, and Roc1b/Roc2 were lethal, and whether Cul-1 and Elongin C showed severe growth defects. When knocked down by *SD-Gal4,* Cul-4 showed a severe wing phenotype, probably via Cdt1degradation (Table 2 and Fig. 2).

**Table 2.** Phenotype of overexpression by *SD-Gal4*.

| Responder (UAS-IR) | Chromosome linkage | Wing phenotype/total flies (%) |
| --- | --- | --- |
| Mcm2 | III | 32.20% (N = 202) |
| Mcm3 | III | Normal (N = 21) |
| Mcm4IR-1 | II | 13.81% (N = 128) |
| Mcm4IR-2 | III | 17.73% (N = 141) |
| Mcm5IR-1 | II | 8.33% (N = 72) |
| Mcm5IR-2 | III | 23.64% (N = 128) |
| Cdc6IR-3 | II | Normal (N = 52) |
| Cdt1IR-1 | II | Lethal (N = 18) |
| Cdt1IR-2 | III | 68.97% (N = 29) |
| Orc4IR-3 | II | 4.12% (N = 120) |
| Orc4IR-2 | III | 3.09% (N = 97) |
| Orc5IR-2 | III | Normal |
| Orc6 | II | 17.31% (N = 103) |
| RPA70IR-1 | II | 95.49% (N = 133) |
| RPA70IR-2 | III | 100% (N = 21) |
| Psf1 IR-2 | III | 24.56 % ( N = 57) |
| Psf2 IR-4 | II | 71.96% (N = 214) |
| Psf2 IR-1 | III | 20.47% (N = 215) |
| Rfc3 IR-9 | II | 100% (N = 158) |
| Pol $\alpha$ 180IR-1 | III | 25.45% (N = 55) |
| Pol $\alpha$ 180IR-3 | III | 50.88% (N = 228) |
| Pol $\alpha$ 50IR-1 | II | 100% (N = 12) |
| Pol $\alpha$ 50IR-2 | III | 100% (N = 146) |
| Pol $\delta$ 125KDa<br>IR-1 | II | 92.86% (N = 84) |
| Pol $\epsilon$ 255IR-1 | II | Normal (N = 153) |
| Pol $\epsilon$ 255IR-2 | III | Normal (N = 180) |
| Cul-4 | II | 86.41 % ( N = 103) |
| Cul-1 | II | Normal (N = 126) |
| dskp-1/SkpA | III | Normal (N = 7) |
| Roc1b/Roc2 | III | Normal (N = 43) |
| Elongin C | III | Normal (N = 160) |
| DP1IR-2 | III | 100% (N = 82) |

\*Red: more than 50% of flies shows wing phenotype.

### The knockdown of E2F and RB by SD Gal4 drivers disturbed wing development

The Rb/E2Fpathway is critical in regulating the initiation of DNA replication (Nevins, 2001). E2F1 and DP mutants, and the knockdown of DP1 by *Act5C-Gal4* and *Tublin-Gal4*, is lethal prior to the adult stage. In 100% of flies, the knockdown of DP1 by *SD-Gal4* resulted in a wing phenotype. These results showed that E2F-DP1 contributes to wing formation.

## Discussion

### The knockdown of DNA replication machinery, SCF complex, and E2F-DP1 by SD Gal4 drivers disturbed wing development

The tissue-specific knockdown of gene expression in *Drosophila* resembled that of the *Cre-loxP* system in mice (Lee and Carthew, 2003; Carthew, 2003). In a cross between a *Gal4* driver line and a line with a UAS small hairpin RNAi transgene construct, the RNAi target mRNA was eliminated temporally and spatially in a tissue-specific manner (Lee and Carthew, 2003; Carthew, 2003). We found that the knockdown of a number of components of the DNA replication machinery caused a rough eye phenotype (Kohzaki et al., 2018a; Kohzaki et al., 2018b). Among them, Mcm10 is likely to function not only during the wing formation and growth phase, but also during differentiation, and in DNA replication.

At the growth/differentiation transition (GDT) point (Jasper, 2002), differentiation signals are expected to enter into chromosomal DNA replication machinery (Kohzaki and Murakami, 2005). Mcm10 may be the endpoint of these wing formation signals. Screening and the use of various mutants will clarify this.

Interactions of DNA replication machinery have been investigated through *in vitro* analysis, such as in *S. cerevisiae* or cell culture systems, but not in the morphogenesis of higher eukaryotes. We revealed these interactions in *Drosophila in vivo* based on phenotype (Basler and Hafen, 1991). This shows that the DNA replication machinery functions as developmental players during development and differentiation. In future, we will extend our analyses to other tissues.

The knockdown of Cul-4, one of the components of the SCF complex, by *SD-Gal4*, showed a severe wing phenotype (Table 2 and Fig. 2). But in the case of dSkip-1/SkpA, Roc1b/Roc2, Cul-1, and Elongin C, we observed no wing phenotype, suggesting that the phenotype of Cul-4 likely results from Cdt1degradation, because the knockdown of Cdt1results in a wing phenotype or lethality, even when knocked down by *SD-Gal4*.

E2F-DP1 is a key player in cell cycle progression, and Buttitta et al. (2007) showed that the crosstalk between E2F and cyclin/Cdk activities was repressed as cells terminally differentiate in *Drosophila* wings. Therefore, the activities of both E2F and the G1 cyclin/Cdks must be simultaneously increased to force these cells to bypass cell cycle exit, or to re-enter the cell cycle after differentiation. In the wing epithelium, additional unknown mechanisms contribute to the downregulation of E2F and G1 cyclin/Cdk activities upon differentiation (Buttitta et al., 2007), partially via Orc1 expression. As a next stage, we must investigate the relationship between E2F functions and terminal differention.

## Acknowledgements

We thank Dr. Masamitsu Yamaguchi (Kyoto Institute of Technology) and his lab for their technical advice. We thank Dr. Tadashi Uemura (Kyoto University), Dr. Kamei (Kyoto Institute of Technology), and their lab for their dedicated support and helpful assistance. This work was partially supported by the Japanese Leukemia Research Fund. H.K. was supported by a KIT VL grant, the Memorial Fund on the 44th Annual Meeting of the Japan Society for Clinical Laboratory Automation and The Motoo Kimura Trust Foundation for the Promotion of Evolutionary Biology.

